# A framework for efficient CRISPRi-mediated silencing of retrotransposons in human pluripotent stem cells

**DOI:** 10.1101/2025.10.24.684366

**Authors:** Anita Adami, Raquel Garza, Fereshteh Dorazehi, Christopher H. Douse, Johan Jakobsson

## Abstract

This protocol describes a workflow for transcriptional silencing of transposable elements (TEs) in human induced pluripotent stem cells (hiPSCs). It illustrates how to design gRNAs to target TE families or unique TE loci and how to validate the efficiency and specificity of large-scale CRISPRi-based silencing using a multiome approach that combines bulk RNA sequencing, CUT&RUN epigenetic profiling, and proteomics. This unique framework optimizes performance and interpretation of *in vitro* functional studies based on transcriptional manipulation of TEs in hiPSC models.

For complete details on the use and execution of this protocol, please refer to Adami *et al*. (2025)^1^.

**Graphical abstract:** 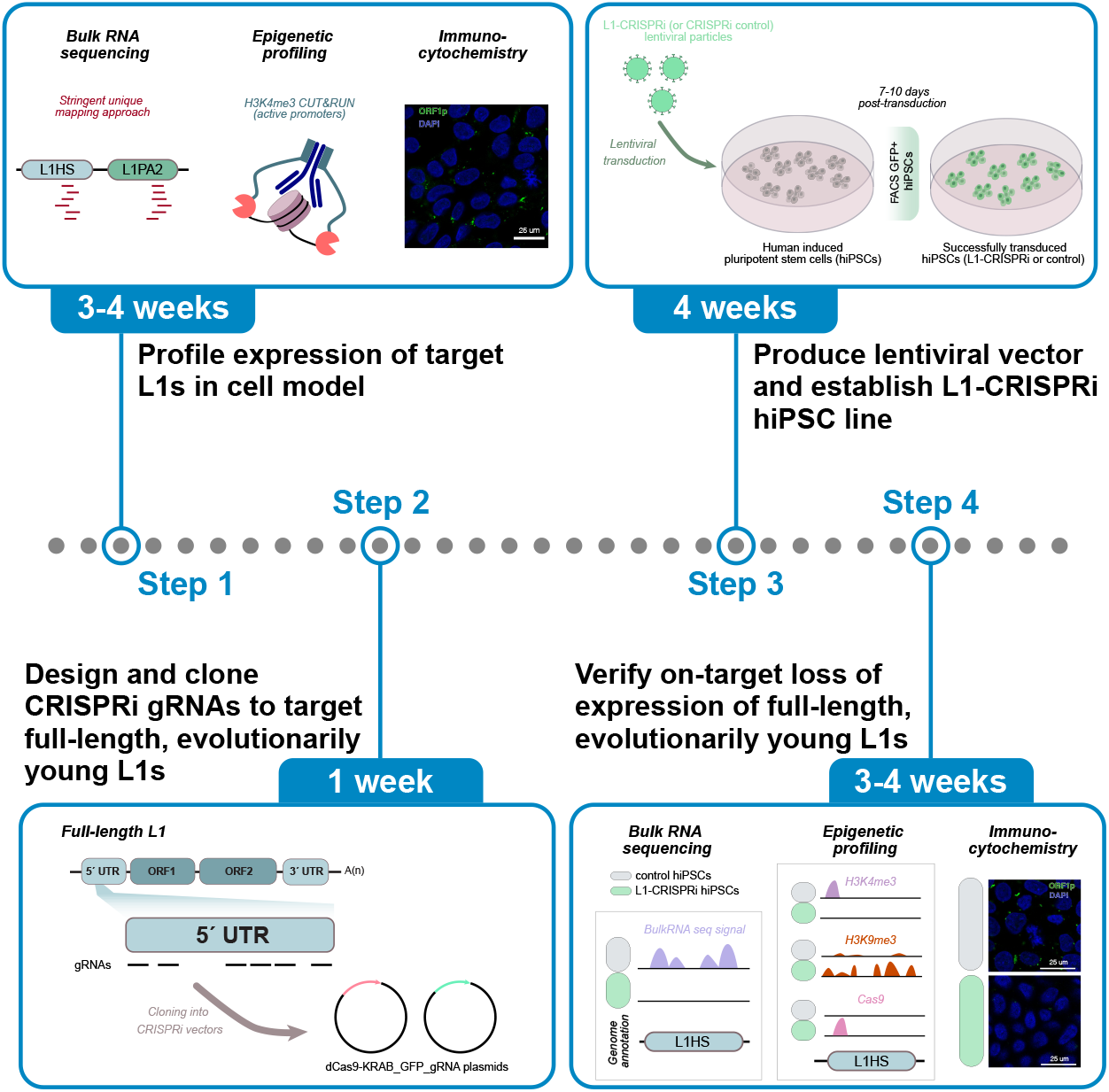

## Before you begin

Transposable elements (TEs) are repetitive sequences that have colonised the human genome throughout evolution and comprise approximately 50% of the human genome^2,3^. Historically viewed as genomic relics, TEs have been severely understudied. Their repetitive nature poses considerable challenges in the investigation of these elements; however, recent advances in sequencing technologies and off-target-minimised CRISPR-based approaches now enable thorough TE investigation at the locus-specific and subfamily levels. Still, downstream on- and off-target analyses of transcriptional manipulations are needed to validate the specificity of these tools and ensure that any observed phenotypic effect in a chosen *in vitro* model is in fact a consequence of the TE manipulation.

The protocol described below provides detailed steps for designing gRNAs to target TE families (multiple copies) using CRISPR inhibition (CRISPRi) and to validate the successful targeting of multiple sequences. Specifically, we present a solid framework to thoroughly assess the specific and efficient silencing of evolutionarily young long interspersed nuclear elements-1 (L1s) (Figure 1) in human induced pluripotent stem cells (hiPSCs). Recent studies have shown that L1s are functionally embedded into transcriptional regulatory networks^1,4,5^ and play a significant role in embryonic development by influencing the chromatin state of their genomic location^6,7^. Thus, it is relevant to investigate L1s in their specific genomic context, which can be achieved, for instance, via their transcriptional manipulation using CRISPRi, as described here.

**Figure 1:**
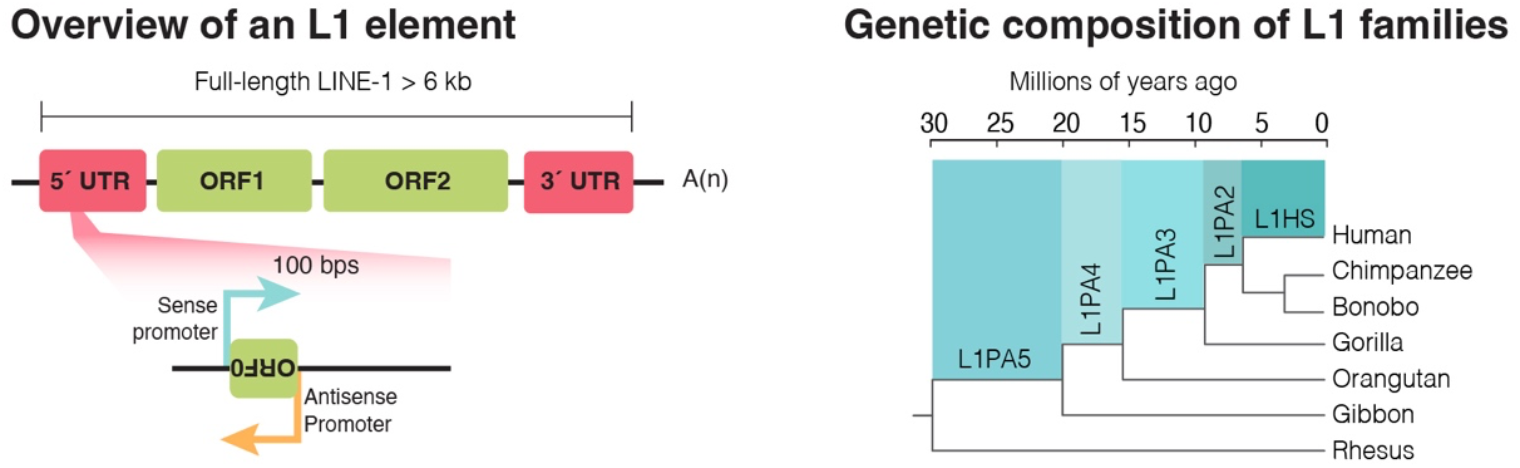
Long interspersed nuclear elements (LINE-1s). Left: Overview of the structure of a full-length, evolutionarily young L1 elements which include an intact 5’ untranslated region (UTR), three open reading frames (ORFs), and a 3’ UTR. Right: Phylogenetic tree showing the evolutionary age of the younger L1 subfamilies.

This protocol can also be used to design gRNAs for CRISPR-based transcriptional manipulation of other TE subfamilies, such as human endogenous retroviruses-K (see also Padmanabhan Nair *et al*., 2022^8^).

We performed the CRISPRi experiments using an all-in-one CRISPRi backbone from Addgene^9^ (RRID: Addgene_71237) into which we cloned the different gRNAs. For the successful application of this protocol, techniques such as molecular cloning, lentiviral production, sequencing infrastructure accessibility, hiPSC culture routines, and bioinformatic expertise must be established in the lab. These techniques are not thoroughly covered here.

### Innovation

This protocol presents a novel, robust framework for targeting repetitive sequences in human pluripotent stem cells (hPSCs) with CRISPR-based transcriptional silencing. We describe a multiomics workflow that uses established molecular and computational techniques to allow for a detailed validation of the specificity and efficiency of the experiment. The novelty lies in the complementary integration of bulk RNA sequencing, unique-mapping analyses for locus-specific target identification and CUT&RUN analysis for detailed epigenetic profiling within a comprehensive, step-by-step pipeline. This framework provides a standardized strategy for designing and applying a TE-targeting CRISPRi system and thoroughly assessing its on- and off-target effects. This systematic approach ensures reproducible and reliable TE transcriptional silencing and establishes an innovative, comprehensive workflow for studying TEs in hPSCs.

### Institutional permissions

The handling of and the data generation from all the human induced pluripotent stem cell lines has been performed in accordance with institutional as well as national regulations.

### TE expression profiling *in vitro*

#### Timing: 3-4 weeks

Before designing and establishing the TE-targeting CRISPRi system, it is essential to profile the expression of the TE family in the chosen *in vitro* model to adequately design the downstream investigations. In this study, we performed an in-depth multiomics analysis including bulk RNA-sequencing, CUT&RUN epigenetic profiling, and proteomics to quantify the expression of evolutionarily young, full-length (> 6kbp) L1s in human induced pluripotent stem cells (hiPSCs).

**CRITICAL:** Profiling the expression of L1s (or the TE family of interest) in the chosen model *before* performing any transcriptional manipulation experiment provides the basis for target selection and ensures resources are not wasted on silencing unexpressed elements. Furthermore, this is key to adequately interpret the transcriptional perturbations introduced with the CRISPRi system and perform a locus-specific analysis of the targeted TEs.

1. Culture hiPSCs and harvest pellets for bulk RNA-seq (stranded libraries, paired-end 2 x 150 bp, poly(A)-selected). A minimum of three technical replicates is generally recommended for appropriate statistical methods. **CRITICAL:** It is important to perform *stranded* RNA sequencing to accurately quantify the transcription of the L1 sequence *in sense* (potentially originating from the element’s promoter), or *in antisense*, (potential read-through signal coming, e.g., from nearby or overlapping protein-coding genes).
2. Analyse bulk RNA-seq data using a unique mapping approach (See also bulk RNA sequencing analysis of L1-CRISPRi hiPSCs within this publication). **CRITICAL:** Apply a unique-mapping strategy to detect locus-specific TE transcription. This ensures the detection of L1s on a single-locus level and avoids any mapping ambiguities caused by the repetitive nature of these elements.
3. Perform H3K4me3 CUT&RUN epigenetic profiling^10^ to identify active promoter markers of evolutionarily young L1s.

**Note:** This is a relevant step to assess whether these young elements are expressed from their own promoter, or they undergo passive transcription initiated from another gene or TE promoter^4,11^. H3K4me3 CUT&RUN profiling is particularly valuable for studying the intrinsic expression of evolutionarily young TEs, whose highly repetitive nature poses significant mappability challenges. Pairing short-read bulkRNA sequencing, which does not overcome the mapping ambiguities of the highly repetitive sequences, with epigenetic profiling allows to assess the promoter activity of the youngest, lowly mappable, elements and address the mappability issues by unequivocally identifying the-mappable-5’ end of evolutionarily young L1s and its surrounding genomic sequence where the mark spreads^11^ (see also: Adami *et al*., 2025^1^ [Figure 2F]).

**Figure 2:**
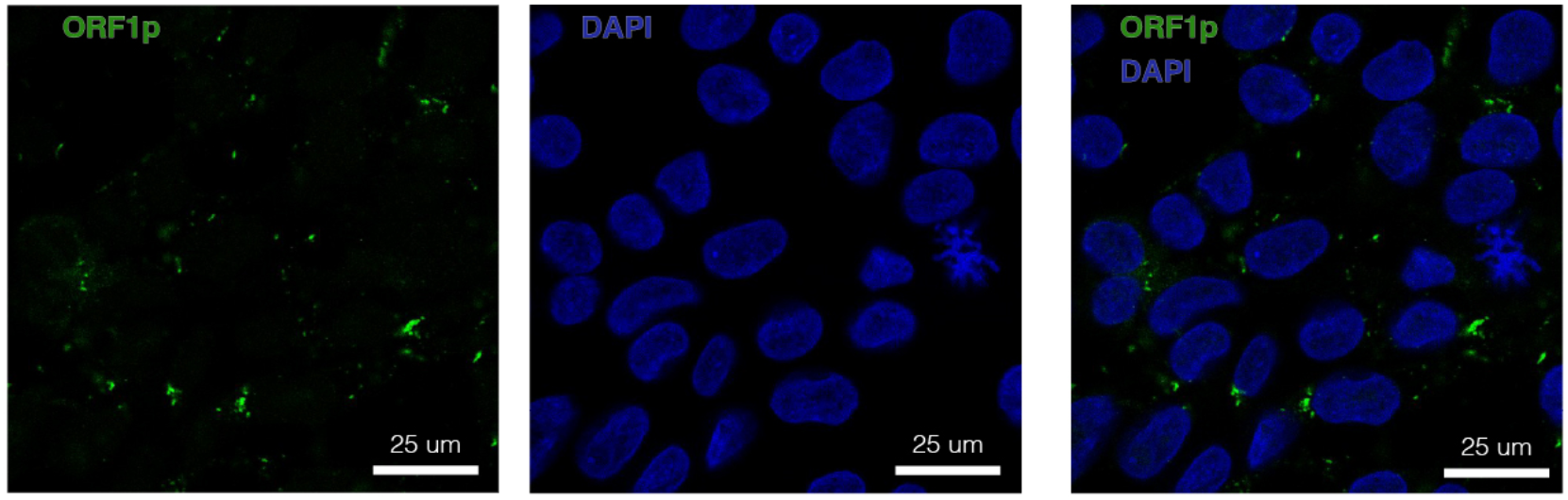
L1-derived ORF1p. Immunocytochemistry of the L1-derived protein ORF1p in human induced pluripotent stem cells. ORF1p is shown in green and a blue nuclear dapi staining is included.

**Note:** If there is no promoter activity from the elements of interest and they are expressed through passive transcription initiated from another gene or TE promoter, it might be relevant to target the promoter that drives the transcription. In other cases, an element that is not actively transcribed might get transcriptionally activated upon a specific treatment, and the experimental design can therefore require the targeting of the newly activated element. However, these are different types of investigation, and they are not further discussed in this publication.

Perform a Western blot and/or an immunostaining (Figure 2) against the L1-derived protein ORF1p to determine whether the expressed L1 elements produce protein.

**CRITICAL:** It is essential that the antibody used is validated and specific to the target. If working with a different subfamily of protein-encoding TEs (e.g. HERV-K elements), this might not be trivial, as many antibodies against TEs can be unspecific.

## Key resources table

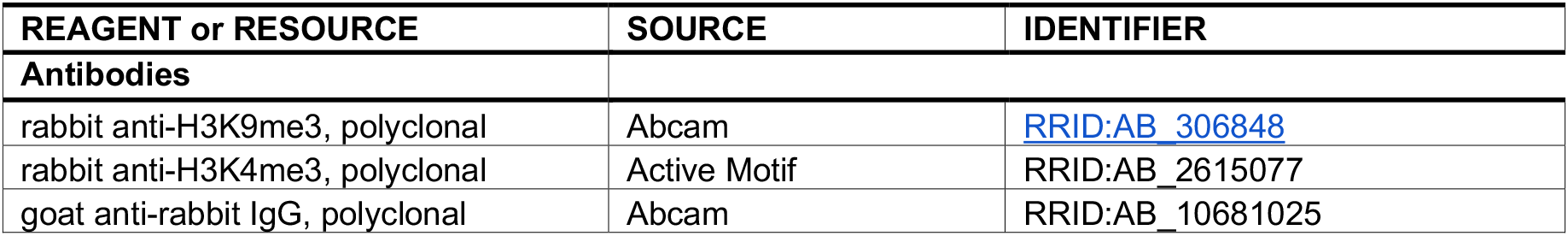
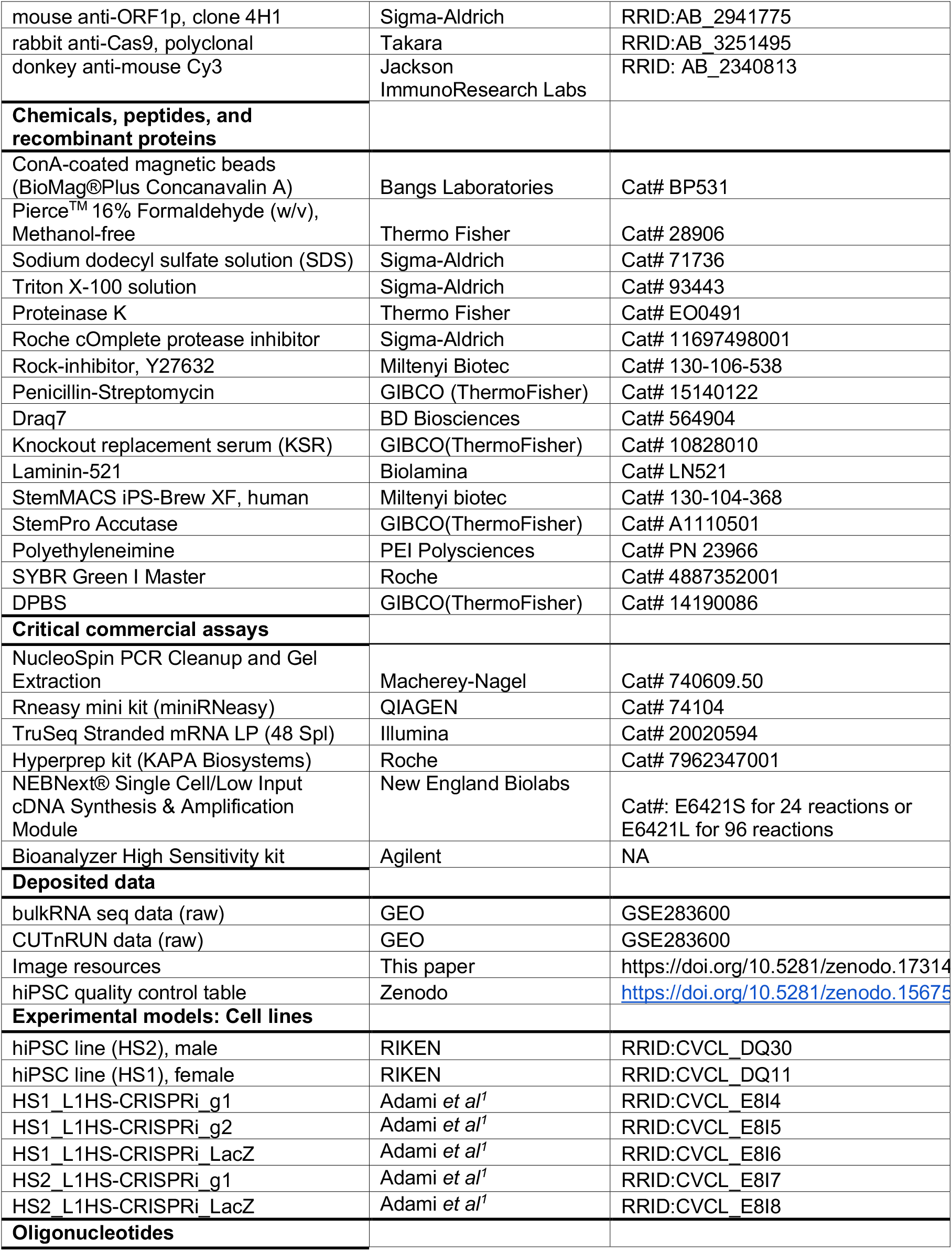
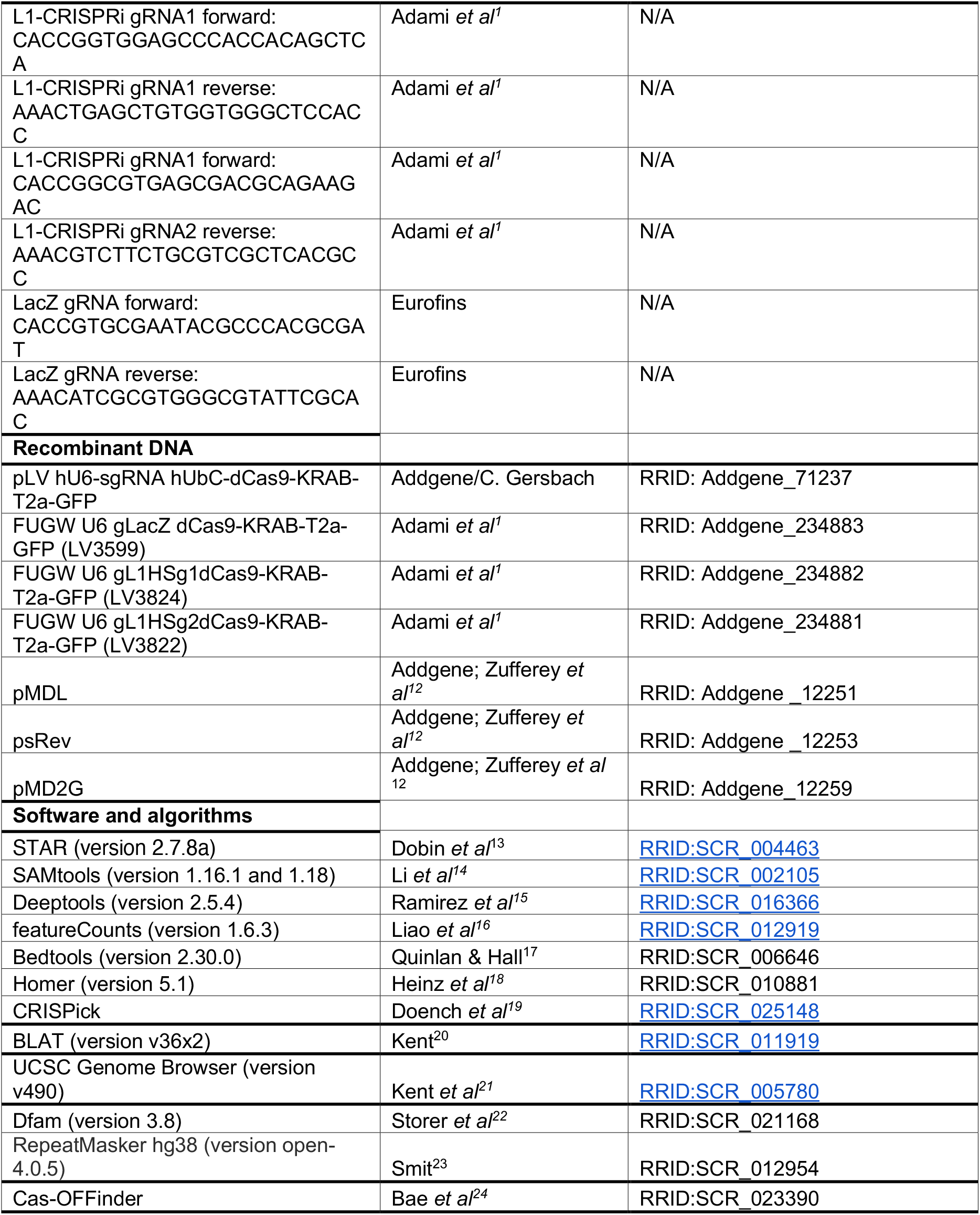

## Step-by-step method details

### TE-targeting gRNA design (TE subfamily level)

Here, we describe the steps to design CRISPRi guide RNAs to target TEs on a subfamily level and accomplish silencing of multiple loci.

#### Timing: 1-2 days

1. Retrieve the consensus sequence for the targeted TE-family as a FASTA file from the Dfam database (RRID:SCR_021168; version 3.8; https://www.dfam.org/home)^22^, which includes an open collection of TE consensus sequences. In our case study, we target evolutionary young L1s (Figure 1) and designed gRNAs against a consensus sequence of the 5’ regulatory region (5’ UTR) obtained from the alignment of the 5’ UTR of human-specific L1HS elements and the hominoid-specific L1PA2 subfamily. This results in the design of gRNAs with the potential to target both L1 subfamilies. **Note:** At this stage, an alternative option is to perform literature research to find already published gRNA designed to target the TE family or subfamily of interest.
2. Design gRNAs against the desired region within the consensus sequence. It is possible to proceed in one of two ways:
  a. ***Option 1*:** Manually design the desired gRNAs.
    i. Search for Protospacer Adjacent Motif (PAM) sequences (NGG for the chosen dCas9-based vector in this study) within the TE consensus sequence.
    ii. Manually design 20 bp-long gRNAs targeting a sequence that is directly adjacent to the identified PAM.

**Optional:** Check potential off-targets with an appropriate webtool, for instance Cas-OFFinder^24^ (RRID:SCR_023390; http://www.rgenome.net/cas-offinder/), for *in silico* predictions. This allows for a quick but comprehensive overview of the designed gRNAs and their predicted target loci and potential off-targets. However, the results when using this type of webtools might be hard to interpret since the designed gRNAs have the potential to target thousands of loci in the genome. Therefore, we recommend proceeding to Step 3 of the protocol, BLAT your gRNAs (TE subfamily level).

**Note:** The PAM sequence is required for the dCas9 machinery to bind the target DNA and needs to be located at the 3’ end of the DNA target sequence of choice and will be directly adjacent to the designed gRNA (Figure 3).

**Figure 3:**
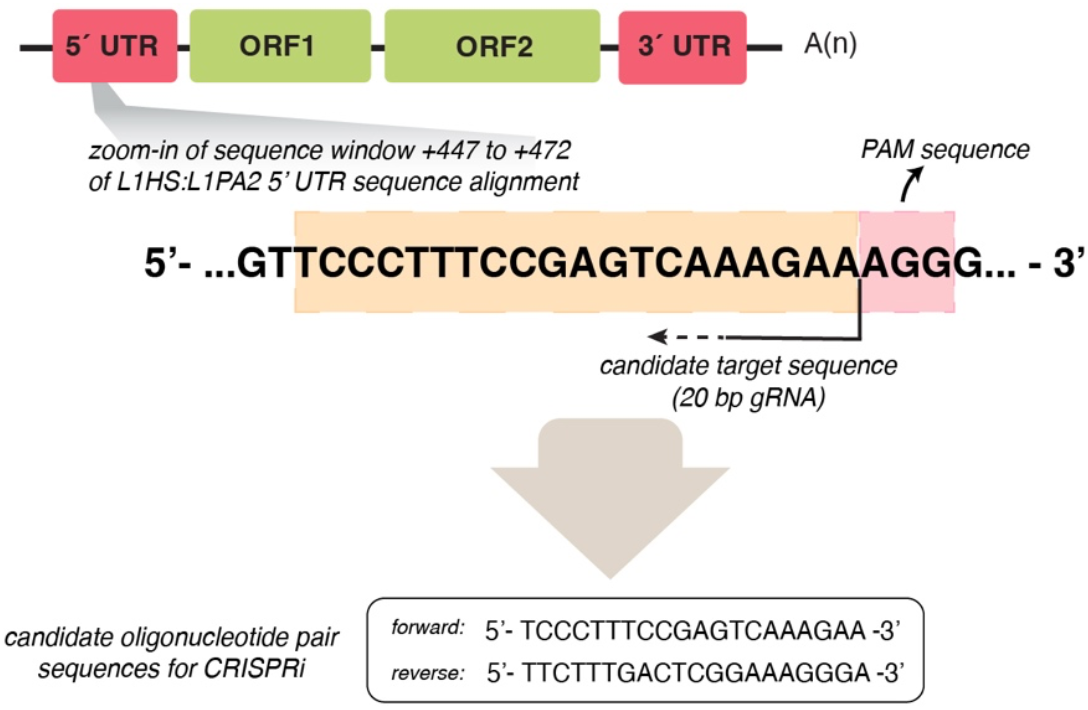
Example of a manually designed gRNA. The designed gRNA (oligonucleotide pair) targets the 5’ UTR of full-length, evolutionarily young L1s.

b. ***Option 2:*** Treat the consensus sequence from Dfam as a single-copy gene.
  i. Input the consensus sequence retrieved from Dfam into a gRNA-designing software such as CRISPick (RRID:SCR_025148; https://portals.broadinstitute.org/gppx/crispick/public)^19^.
  ii. Define the conditions of your experiment: Reference genome Human GRCh38 (or the more recent annotation T2T^3^ if it is implemented in the chosen gRNA design software/tool); Mechanism-CRISPRi; Enzyme-SpyoCas9.
  iii. Upload the FASTA file retrieved from Dfam with the desired consensus sequence.
  iv. Inspect the resulting.txt file on Excel and pick the desired candidate gRNAs based on scoring and minimum predicted off-target effect.

**Note:** It is still relevant to consider gRNAs that present the minimum predicted off-target effect, as this means that the target sequence does not have many mismatches with the designed gRNA. This is an important point since the sequence of evolutionarily young TEs from the same, or a close, subfamily (e.g. L1HS and/or L1PA2 elements) does not diverge significantly-hence the low mappability of these elements.

For more details on how to use the CRISPick tool, see also: How CRISPick Works and Johansson *et al*. (2022)^25^.

**Note:** Given the repetitive nature of the elements, each gRNA potentially targets different elements within the chosen subfamily. It is therefore preferable to select multiple gRNAs (at least 3-4) before proceeding with the next step to ensure a robust understanding of potential targets, off-targets, and downstream functional consequences of the CRISPR-based manipulation.

### Alternative: TE-targeting gRNA design (single-locus level)

This step presents an alternative on how to design CRISPRi gRNAs that target a single transposable element establishing a silencing on a single-locus level.

#### Timing: 1-2 days

Depending on the research question, it might be of interest to target one single L1 element on a locus-specific level. To do so, gRNAs can be designed as ***Option 2*** above and the single locus-targeting gRNA can be run through the BLAT tool (RRID:SCR_011919; version v36×2) on the USCS Genome Browser (RRID_SCR_005780; version v490) making sure that the designed gRNA only targets one single locus in the genome. Depending on the target element, it can be relevant to consider the sequence right upstream of the element itself: its unique genomic location will minimize off-target effects of the designed gRNAs.

**Note:** When performing a TE-targeting on a single-locus level, it is also crucial to design multiple (e.g. 2 or 3) gRNAs for the CRISPRi experiments. This ensures that any phenotypical effect observed following the transcriptional silencing of the targeted elements is not to be attributed to off-targets, since different gRNAs will also have different potential off-targets.

### BLAT your gRNAs (TE subfamily level)

The following steps illustrate how to verify *in silico* that the designed gRNAs target the TE subfamily (or subfamilies) of interest.

#### Timing: 1 day

**Note:** Given that the designed gRNAs will target multiple loci in the genome, it is important to predict *in silico* the targets for each candidate gRNA to proceed with those that present the maximum on-target effects and minimum off-targets.

3. BLAT the sequence either using the BLAT tool on the UCSC Genome Browser, or using the command-line version available for download from the UCSC servers. If using the command-line version, define the following parameters:
  a. Set a small stepSize (1-5 bps). **Note:** Step sizes define the frequency at which to query the CRISPR guide into the reference genome (target sequence). For instance, in a target sequence of length 100 bp and a query of 10 bp, a stepSize= 10 will query the sequence to the target every 10 bps, resulting in 10 comparisons. A stepSize= 5 will query the sequence to the target every 5bps, resulting in 20 comparisons. **CRITICAL:** The default stepSize in BLAT is optimized for speed, not sensitivity. Thus, lowering the stepSize (e.g. to 1-5 bps) increases the chances of detecting interrupted alignments (non-perfect matches), which is especially relevant for old TE subfamilies (e.g., ERVs) where several mutations have accumulated throughout evolution, or subfamilies prone to carry polymorphisms within their sequence (e.g., SVAs).
  b. Set a large repMatch. **Note:** This parameter defines the number of times the query (gRNA) is allowed to appear in the target sequence (reference genome). For TEs, this number should not be smaller than the size of the subfamily in question (minimum number of elements found in the chosen TE subfamily-e.g. L1HS: 1642 elements, which can be rounded up to include closely related sequences).
  c. Set a sensitive minScore. **Note:** This threshold defines the minimum alignment quality for a match to be reported, and it will influence the number of hits you get reported by BLAT. The score is calculated based on the number of unique matches in the genome (+1), the number of multimapping matches (+0.5), mismatches (-1), and gaps (-1). Note that the number of multimapping matches are counted as half but are likely to inflate TE scores. For the purposes of TE analyses, where even weaker alignments are relevant, we recommend lowering the default minScore to evaluate partial matches to the target sequence. It is however important to know that setting a lower minScore can potentially lead to false positives. To address this, see Problem 1: Clear false positive hits in the BLAT analysis in the Troubleshooting section.

Taken all these parameters together, here is an example of how a BLAT command could look like:

~~~
blat -stepSize=1 -repMatch=2000 -minScore=15./genome.fa./guide1.fa./guide1.blat.out
~~~

**Note:** If using the online BLAT tool on the UCSC Genome Browser, stepSize, repMatch, and minScore parameters cannot be manually defined. This can limit the *in silico* predictions as the search might not be sensitive enough to pick up partial matches.

4. To interpret the BLAT results, intersect the coordinates using bedtools intersect. Parameter -a (file “A”) being the genomic coordinates of the BLAT results. Flag -wo to report only entries in file “A” with an overlap in file “B”. File “B” being one of the following:
  a. **RepeatMasker annotation:** RepeatMasker^23^ (RRID:SCR_012954; version open-4.0.5) is a software that screens a genomic sequence (e.g., the human genome) to annotate TEs and low complexity regions. To download the RepeatMasker human annotation, visit the following link: https://www.repeatmasker.org/species/hg.html. Both the annotation file and the BLAT results should be converted to BED format (https://genome.ucsc.edu/FAQ/FAQformat.html#format1) to be used with bedtools intersect and evaluate the specificity to a particular subfamily or element. **Note:** This is especially relevant over composite elements, where CRISPR guides could simultaneously target several subfamilies due to their evolutionary history and sequence composition. For instance, SVA-targeting guides could also target Alus and/or ERVs, since SVAs share portions of their sequence with these other TE families.
  b. **Genes’ exons annotation:** To evaluate potential targeting of gene sequences (potential off-targets). **CRITICAL:** This should be performed over exons coordinates rather than genes coordinates, to avoid intersecting with potential intronic on-target TE sequences (covered in the next step).
  c. **Genes’ introns annotation:** To annotate intronic targets. **Note:** This step complements the previous two, by classifying targeted elements either as intergenic or intragenic.

Taken the parameters, an example of a BEDtools intersect command could look like:

~~~
bedtools intersect -a guide1.blat.bed -b gencode.gtf -wo > guide1.blat.intersect.gencode.out
~~~

**Note:** After analyzing the BLAT results, 2-6 gRNAs will likely stand out as good candidates for the formulated research question (e.g. they exclusively, or mostly, target the TE subfamilies of interest). These 20-nucleotides sequences will be the gRNAs chosen to proceed with the experimental work described below.

### Clone the gRNAs and produce silencing lentiviral vectors

Here, we present an overview of the steps that lead to the production of the CRISPRi lentiviral vectors.

#### Timing: 3 weeks

5. Order the oligos containing the designed and chosen 20 bp gRNA and the correct flanking restriction sites (BsmBI in this study) and clone them in the CRISPRi plasmid backbone (we used RRID: Addgene_71237).

Forward oligo: 5’ **CACCG**…20bp gRNA… 3’

Reverse oligo: 5’ **AAAC**…20 bp gRNA…**C** 3’

**Note:** Clone one gRNA per plasmid.

**CRITICAL:** It is crucial to include a non-targeting gRNA (e.g. against the bacterial gene *lacZ*, not present in the human genome, or a scrambled sequence) as a negative control for all the CRISPRi experiments. Plasmids including this gRNA should be produced in parallel with the other CRISPRi plasmids. See the Key Resource Table included in the paper for the exact sequences used in this study.

6. Produce lentiviral vector from the newly cloned plasmids. Lentiviruses for the L1-targeting and non-targeting control vector were produced according to the third generation lentiviral production protocol by Zufferey *et al* (1997)^12^. Virus titration was performed via qPCR of genomic DNA from serial dilution transductions. Titers no lower than 10^8^ TU/ml were obtained and used in the experiments.

### L1 CRISPRi in hiPSCs via lentiviral transduction

The following steps describe how to perform the lentiviral transduction to establish the desired CRISPRi hiPSC lines.

#### Timing: 2 weeks

**Note:** Firstly, the L1-CRISPRi hiPSCs line (and the correspondent negative control) needs to be successfully established via lentiviral transduction. For more details on hiPSCs culture, see also Grassi *et al*. (2020)^26^, Johansson *et al*. (2022)^27^, Adami *et al*. (2025)^*1*^. The quality controls performed on the hiPSCs used in the original publication can be found at the following link: https://doi.org/10.5281/zenodo.15675716.

7. **Establish the L1-targeting and the non-targeting control hiPSCs lines**.
  a. Culture hiPSCs on Lam521-coated wells and change media every day using iPS-Brew or an equivalent iPSCs media. Passage cells every 3-4 days using Accutase, seeding 100 000-200 000 cells/well in a 12-wells plate with 900 μl of media supplemented with 10 μM of Rock inhibitor.
  b. After expanding the hiPSCs (at least one passage after thawing), transduce cells:
    i. Seed 250 000 cells/well in 900 μl of media in a 12-wells plate (at least 1 well/condition, including the negative control).
    ii. Within 2-5 minutes after seeding (before the cells start attaching to the well), add the equivalent of multiplicity of infection (MOI) 10 of the CRISPRi lentiviral vector to each well. Add **only one virus** per well.

##### Practical example

The virus titer, measured via genomic PCR (gPCR), is 1.63χ10^9^ TU/ml, and the chosen MOI (average number of viral vector particles per cell) for the experiment is 10. In a 12-wells plate, seed 250 000 cells/well for the transduction. To calculate how much volume of the lentiviral preparation is needed to achieve the desired MOI, apply the following formula:

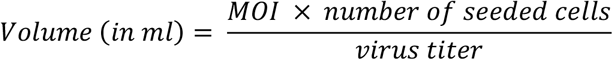

To calculate the amount of virus needed in μl, multiply the result times 1000:

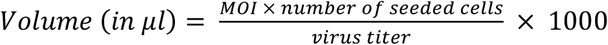

Therefore, the calculations for the given example are:

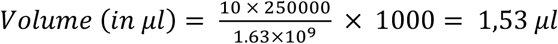

**Optional:** Since the resulting volumes might be very small to pipette (> 1 μl), a 1:10 dilution (in iPS-Brew or equivalent iPSCs media) of the viral preparation can be prepared to pipette larger volumes.

**Note:** Adding one lentiviral vector (*i*.*e*. one gRNA) per well avoids mixed silencing effects. Comparable effects observed using different gRNAs ensure a more robust interpretation of the phenotypical consequences of the TE silencing.

c. Immediately after adding the virus, add double media (e.g. 900 μl× 2= 1800 μl in each well of a 12-wells plate) and supplement with 10 μM Rock inhibitor.
d. Change media and supplement with Rock inhibitor until the first split, ideally 3 days post-transduction. At this point, the cells should have recovered from the lentiviral transduction, a good indication of when to proceed with the first split.
e. By day 7-10 after transduction, the CRISPRi system has been constitutively established and the hiPSCs can be selected. Using fluorescence-activated cell sorting (FACS), GFP+ hiPSCs can be sorted. The GFP+ cell population corresponds to the successfully transduced cells and can be collected for bulk RNA sequencing and cryopreservation:
  i. Sort 100 000-200 000 GFP+ cells via FACS.
  ii. Pellet at 500 x *g* for 10 minutes.
  iii. Process cells for various downstream purposes.
  iv. Remove supernatant and snap freeze for **bulk RNA sequencing** (stranded libraries, paired end 2 x 150 bp).
  v. Collect 100 000-200 000 cells/condition, replate them on Lam521-coated wells (12-wells plate) and expand for **cryopreservation and/or downstream experiments**.

If excessive cell death is observed following the lentiviral transduction, see Problem 2: Extensive cell death across all conditions after transduction in the Troubleshooting section.

If not enough GFP+ cells could be recovered from FACS, see Problem 3: Not enough GFP+ cells can be recovered after transduction in the Troubleshooting section.

**Note:** Selection of GFP+ cells is crucial to ensure only cells with silencing of young L1s in downstream assays. If the cells are cryopreserved, it is preferrable to resort the GFP+ population prior to any downstream application (e.g. differentiation of CRISPRi hiPSCs into unguided cerebral organoids, as we report in Adami *et al*. (2025)^*1*^).

**Note:** All the steps described within this section (L1 CRISPRi in hiPSCs via lentiviral transduction) apply to both the multi-loci silencing of evolutionarily young L1s and the single-locus CRISPRi experiments.

### Bulk RNA sequencing analysis of L1-CRISPRi hiPSCs

The following step illustrates how to leverage bulk RNA sequencing data to check the CRISPRi silencing.

#### Timing: 1 week

**Note:** Because TE targeting is often multi-loci-targeting, either by design or not, methods like qRT-PCR lack the specificity to analyse transcriptional changes on a locus-specific level. Thus, a bulk RNA sequencing analysis using a stringent unique-mapping approach is crucial to comprehensively assess the consequences of CRISPRi on evolutionarily young L1s. This step is key for any newly designed gRNA, as the *in silico* prediction of the targeted loci might not fully predict experimentally observed effects.

8. Map reads using STAR aligner (we used version 2.7.8a) with the latest genome index and GENCODE annotation as guide GTF (--sjdbGTFfile):
  a. Allow a single mapping locus (--outFilterMultimapNmax 1)
  b. Set a ratio of mismatches to the mapped length of 0.03 (--outFilterMismatchNoverLmax)

**Optional:** It may be informative to determine the sense and antisense transcription originating from evolutionarily young L1s, which harbor a bidirectional promoter in their 5’ UTR (Figure 1). One approach is to use alignment files (BAM or SAM) to generate strand-specific bigwig files using the bamCoverage function of deeptools (we used version 2.4.5). These files can then be inputted into computeMatrix and plotHeatmap deeptools functions to produce heatmaps for L1 subfamilies of interest^4^. For a more quantitative approach, alignment files can be split by strand based on SAM flags using samtools view (we used version 1.16.1), followed by generating count matrices for the evolutionary young L1 elements. This allows assessment of in-sense and antisense transcription over the elements. Additionally, one could also define custom GTF entries for regions upstream of the L1 to capture L1-ORF0-derived transcription (we defined these regions as 500bps upstream the L1 TSS in the opposite direction).

**Note:** This first analysis is sufficient for an initial screening of the CRISPRi gRNAs designed. Testing several multi-loci L1-targeting gRNAs will allow to select the best gRNAs based on specificity and efficiency. Depending on the research question, one can proceed to the next step with 1-3 gRNAs of choice.

### H3K4me3, H3K9me3, and dCas9 CUT&RUN of L1-CRISPRi hiPSCs

Here, we describe the workflow to verify the on-target specificity of the designed CRISPRi gRNAs using CUT&RUN epigenetic profiling.

#### Timing: 2 days (CUT&RUN protocol) + 1 week (CUT&RUN analysis)

**Note**: The repetitive nature of evolutionarily young L1s makes it essential to complement bulk RNA sequencing data (as the youngest elements suffer very low mappability) with techniques such as CUT&RUN epigenetic profiling ^10,28^ to identify transcription (and loss of transcription upon CRISPRi) from active L1 promoters using H3K4me3 CUT&RUN. In addition, H3K9me3 and dCas9 CUT&RUN on the same cells allows a deeper investigation of both on-target and off-target effects of the targeted L1 elements.

We have shared detailed protocols for both CUT&RUN experimental workflows on protocols.io and they can be found at the following links: dx.doi.org/10.17504/protocols.io.36wgqdb83vk5/v1 (histone markers CUT&RUN) and dx.doi.org/10.17504/protocols.io.bp2l6dr5rvqe/v1 (Cas9 CUT&RUN).

9. Perform H3K4me3 and H3K9me3 CUT&RUN epigenetic profiling of L1-CRISPRi hiPSCs (including the non-targeting control cells). **Note:** H3K9me3 is a histone marker found at heterochromatic sites. Performing H3K9me3 epigenetic profiling in L1-CRISPRi hiPSCs (vs control cells) allows for the assessment of H3K9me3 spread around the dCas9 binding site at targeted L1 promoters. This analysis is essential to ensure that the CRISPRi system, which deposits H3K9me3 to silence the targeted loci, is specifically targeting the promoters of evolutionarily young L1s but not nearby gene promoters. However, in cases where the L1 promoter acts as a promoter of an alternative transcriptional isoform of a nearby gene, both the L1 element and the gene might be simultaneously silenced by CRISPRi. By assessing targeting specificity in this way, one can evaluate if downstream phenotypic effects (e.g. transcriptional changes) are a direct consequence of the L1 silencing or off-targets. **Note:** H3K4me3 is an active chromatin mark that can be used for both identifying transcriptionally active L1 elements and measuring on-target efficiency of the CRISPRi system by ensuring the loss of this mark over targeted evolutionary young L1 elements.
10. Perform Cas9 cross-linking CUT&RUN epigenetic profiling of L1-CRISPRi hiPSCs and control cell line.

**CRITICAL:** Even though most targets such as H3K9me3 and H3K4me3 work well under native condition when profiling by CUT&RUN, we have found that dCas9, along with other chromatin-binding proteins, benefit from a light cross-linking condition to improve signal recovery. To this end, cells should be cross-linked using a 0.1% formaldehyde solution (for 1 min) prior to chromatin digestion and release. More details on this protocol can also be found in the original research paper Adami *et al*. (2025)^1^.

**CRITICAL:** We recommend starting with light cross-linking but if that is not sufficient, a moderate cross-linking using 1% formaldehyde solution (for 1 min) could be beneficial. It may be necessary to optimize this step for a new cell type. It is possible that cross-linking compromises the DNA yield. To minimize this issue, see Problem 4: Cross-linking CUT&RUN compromises DNA yield in the Troubleshooting section.

**Note:** A cross-linking CUT&RUN requires changes in wash condition such as addition of 0.05% SDS and 1% Triton X-100 to improve cell permeability and increase yield. Cross-linking must be reversed by adding 0.09% SDS and 0.22 mg/ml proteinase K to the supernatant containing CUT&RUN-digested DNA followed by an overnight incubation at 55 °C.

**CRITICAL:** As formaldehyde is classified as a group 1 human carcinogen, it is crucial to handle it according to relevant safety guidelines, which include the use of a fume hood as well as wearing gloves, eye protection, and a lab coat whenever handling this hazardous chemical.

11. Analyse the CUT&RUN data.
  a. Map reads using Bowtie2 aligner (we used version 2.4.4) with the latest genome index:
    i. For paired-end libraries, use the –no-mixed flag to retain only alignments in which both mates of a read pair successfully mapped.
    ii. Filter out discordant alignments (where mates align separately) of the two pairs using –no-discordant.
    iii. Retain alignments with insert sizes from 10 bps (-I) to 700 bps (-X), corresponding to the distance spanned between both mates of a paired-end read.
  b. Filter out multi mapping by setting the minimum mapping quality score to 10 using samtools view (-q 10).
  c. To visualize the data on genome browsers, use the alignment files (BAM or SAM) to generate bigwig files using the bamCoverage function of deeptools (we used version 2.4.5).

**Optional:** Bigwig files can be input into computeMatrix and plotHeatmap deeptools functions to produce heatmaps for TE subfamilies of interest (see also Garza *et al*. (2023)^4^ for further details).

**Optional:** Peak calling using tools such as HOMER (we used version 5.1) for the Cas9, H3K4me3, and H3K9me3 CUT&RUN datasets can be used to create a curated list of targets displaying the expected effects of the CRISPR targeting, including binding of dCas9, loss of H3K4me3, and gain of H3K9me3. Functional annotation of these sites can be further performed using bedtools intersect to the Repeatmasker, exon, and gene annotations.

## Expected outcomes

The expected outcome for this protocol is the generation of human pluripotent stem cell lines where specific and efficient silencing of evolutionarily young, full-length L1s is established via CRISPRi. In these hiPSCs, a dCas9-KRAB-L1-targeting vector (or control dCas9-KRAB vector) is stably expressed, leading to a multi-loci transcriptional silencing of the targeted subfamily of transposable elements (Figure 4A and 4B). A similar outcome, but with the establishment of constitutive silencing of one single TE, is achieved if the gRNAs have been designed, and properly tested, to target a single locus.

**Figure 4:**
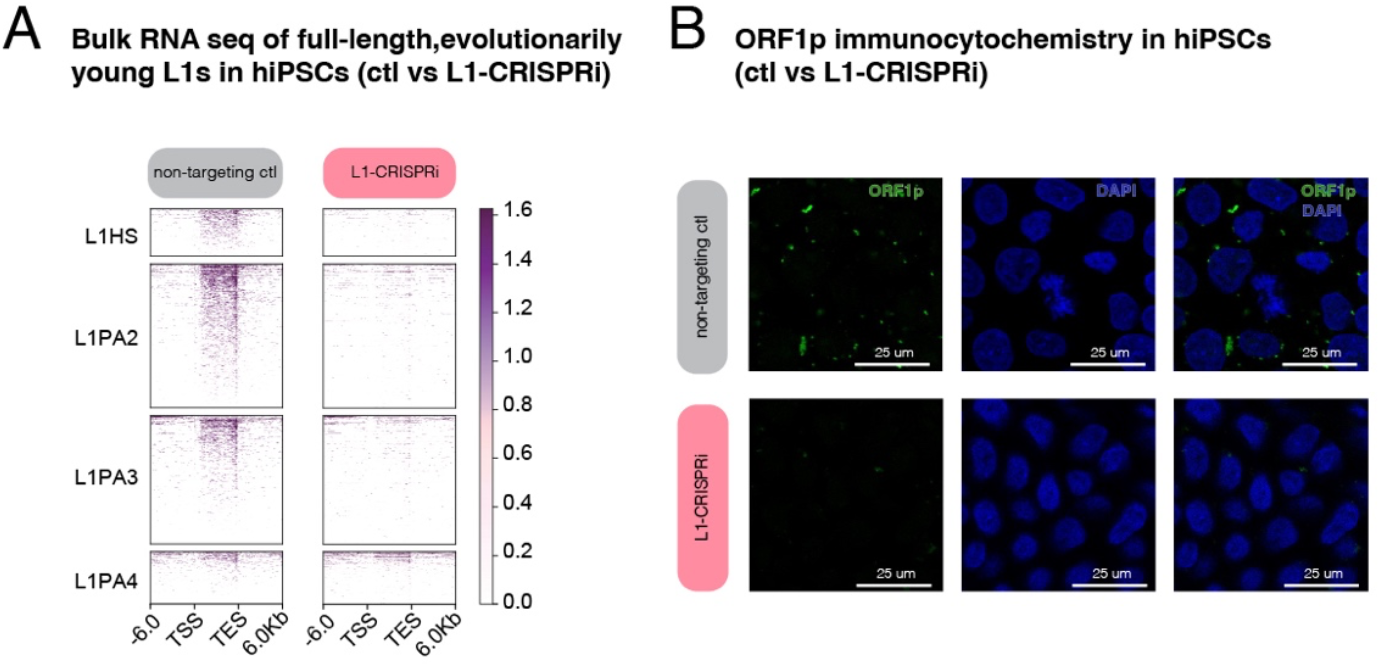
CRISPRi-silencing of evolutionarily young, full-length L1s. A) Deeptool plot showing bulkRNA sequencing analysis (stringent unique mapping) of L1 loci targeted with L1-CRISPRi. The non-targeting control (ctl) is shown on the left, and the L1-CRISPRi on the right. B) Immunocytochemistry of ORF1p in control (non-targeting ctl) vs L1-CRISPRi hiPSCs. ORF1p is shown in green, and dapi nuclear staining in blue.

The generated L1-CRISPRi hiPSC lines and data can be used for downstream analyses (e.g. differential gene expression analysis) to assess any functional consequences that the manipulation of the targeted L1 elements has on the cells. The hiPSCs can be differentiated into other hiPSC-derived *in vitro* models to investigate more specific research questions. For instance, we differentiated our newly generated L1-CRISPRi hiPSCs into unguided cerebral organoids to investigate the consequences of evolutionarily young, full-length L1 silencing in human brain development, leveraging this *in vitro* model to describe the *cis-*regulatory role of these TEs on protein-coding genes and long non-coding RNAs in early human brain development^1^.

## Limitations

This protocol describes how to target evolutionarily young, full-length L1s that have an intact promoter via efficient on-target CRISPRi. One limitation is the use of one single vector per CRISPRi hiPSC line, which makes it quite labor-intensive in terms of cloning, lentiviral vector production, and cell culturing. A CRISPRi vector in which multiple gRNAs can be cloned could be used instead to reduce the production and experimental costs^29,30^.

The presented framework can also be applied for the silencing of other TEs subfamilies. However, the limited mappability of some TE subfamilies (e.g. SVAs) increases the challenges with gRNA design and downstream transcriptomic analyses, including on- and off-target analyses. Therefore, additional considerations for the gRNA design and the on- and off-target analyses might be required.

## Troubleshooting

### Problem 1: Clear false positive hits in the BLAT analysis (Step 3)

We have recommended setting a low minScore in the BLAT analysis to be reported even on weak or partial matches of the guide RNA sequences and allow the identification of all potential TE targets. However, if the minScore parameter is set very low, it can lead to BLAT reporting false positives, i.e., sequences which are unlikely to be true, biologically plausible, CRISPR targets because of the very weak match with the target DNA sequence.

### Potential solution

Post filtering of the false positives can be helpful. Important variables to explore from the BLAT results are the alignment length and percent of identity. Specifically, CRISPR gRNAs are generally 20 bp-long, and if a target sequence matches fewer than 15-16 bases of the gRNA loaded on the Cas9 machinery, it is highly unlikely for the CRISPR system to bind the target DNA. Therefore, BLAT hits with a sequence match of less than 15-16 bases and a mismatch of more than 4-5 bases (out of the 20 bp length of the gRNA) are likely false positives that can be filtered out.

### Problem 2: Extensive cell death across all conditions after transduction (Step 7)

If cell death is observed across all conditions (including the non-targeting control cell line) after lentiviral transduction, this might be a consequence of a viral load too high for the transduced cells to sustain or an impure viral preparation that leads to cytotoxicity.

**Note:** Cell death differences across conditions can be a phenotypical consequence of the specific experiment, but if the control cells also die and do not recover after 2-3 days post transduction, that is linked to the toxicity of the viral transduction itself.

### Potential solution

To solve the problem of a too high viral load, one can try to use lower MOIs and test a few (Figure 5), especially if testing a lentiviral vector for the first time. If the transduction with highest MOI presents extensive cell death across conditions and cannot be maintained to obtain enough transduced cells, then the lower MOIs can potentially survive better to allow for selection by FACS (e.g. GFP+ cell population), and testing for transcriptional silencing.

**Figure 5:**
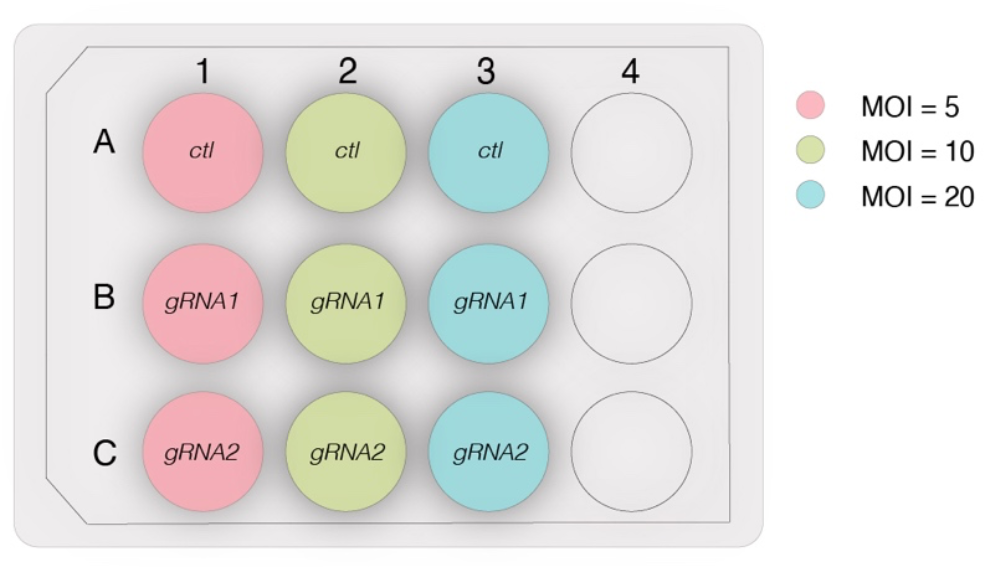
Schematic of a 12-well culture plate with examples of different MOIs to test the lentiviral vector. Ctl= control.

If the problem lies in an impure viral preparation, where contaminants such as cell debris and endotoxins are found in the viral suspension, an optimisation of the plasmid purification and virus production workflow is necessary. It is important to work with high-quality and endotoxin-free plasmid DNA as well as adopt proper filtering and centrifugation techniques when collecting the packaged viral particles.

### Problem 3: Not enough GFP+ cells can be recovered after transduction (Step 7)

If a low percentage of GFP+ cells is recovered during FACS, this could be due to a low virus titre or low efficiency of the lentiviral transduction.

### Potential solution

A high titre of the lentiviral vector is essential to achieve high transduction efficiency, which can be hindered by the presence of unpackaged/free RNA molecules and non-infectious particles in the viral preparation. This leads to a low effective MOI of the viral preparation, with a drastic reduction in transduction and integration efficiency. Optimizing the lentiviral concentration and harvesting methods are essential to obtain high quality and highly concentrated viral particles.

A low transduction efficiency can also occur when the cells are too confluent at the time of transduction. This can lead to uneven transduction across the cells, as well as to a reduction in their metabolic activity, which can prevent optimal transduction. To avoid this, it is crucial to transduce cells which are not overly confluent. An optimal confluency is generally ∼70%, but this can vary (and must therefore be optimized) in different cell types. Transducing cells that have been split no more than twice after thawing can also improve the transduction efficiency.

### Problem 4: Cross-linking CUT&RUN compromises DNA yield (Step 10)

Adding a cross-linking step could be essential when performing CUT&RUN on reversibly bound proteins such as, in this study, dCas9. However, cross-linking might lead to lower DNA recovery.

### Potential solution

To maximize the DNA recovery, it is crucial to handle the sample with extreme care during the cross-linked steps. Loss of DNA could also occur during the washing steps, and to prevent this, it is possible to slightly increase the centrifugation force (e.g. up to 1200 x g). Finally, a higher number of cells as the initial sample input can improve DNA recovery (e.g. 1 000 000 cells per condition).

## Resource availability

### Lead contact

Requests for further information, resources, and reagents should be directed to and will be fulfilled by the lead contact, Johan Jakobsson.

### Technical contact

Technical questions on executing this protocol should be directed to and will be answered by the technical contact, Johan Jakobsson.

### Materials availability

This study did not generate new unique reagents. Plasmids and cell lines generated in relation to the original research study (Adami *et al*., 2025^1^) have been registered on Addgene or Cellosaurus, respectively, are reported in the Key Resource Table included in this publication and are available upon request from the lead contact.

### Data and code availability

No original data or code was generated in this publication.

All the data or code presented in this work have been published in the original research paper Adami *et al*., 2025^1^. The sequencing data discussed in the publication have also been previously deposited at the GEO superseries GEO: GSE283600. The code in relation to the original research publication has been deposited on GitHub and is publicly available at git@github.com:Molecular-Neurogenetics/L1CRISPR_Adami_2025.git (DOI: 10.5281/zenodo.15638389).

The image data presented in this publication have been deposited to Zenodo at doi: 10.5281/zenodo.17314543.

Data, code, protocols, and key lab materials used in this study are listed in the Key Resource Table available within this article and deposited to Zenodo at doi: 10.5281/zenodo.17435053. An earlier version of this article was posted to bioRxiv at https://doi.org/10.1101/2025.10.24.684366.

## Acknowledgments

This study was partially funded by the joint efforts of the Michael J. Fox Foundation for Parkinson’s Research (MJFF) and the Aligning Science Across Parkinson’s (ASAP) initiative. MJFF administers the grants ASAP-000520, ASAP-024296, and ASAP-025170 on behalf of ASAP and itself. The work was also supported by grants from the Swedish Research Council (2022-02673 to J.J. and 2021-03494 to C.H.D.), the Swedish Brain Foundation (FO2023-0232 to J.J.), Cancerfonden (222185Pj to J.J.), Barncancerfonden (PR2023-0099 to J.J.), the Swedish Society for Medical Research (S19-0100 to C.H.D.), and the Swedish Government Initiative for Strategic Research Areas (MultiPark & StemTherapy).

## Author contributions

Conceptualization by A.A., R.G., and J.J.; experimental work was performed by A.A. and F.D.; R.G. executed the bioinformatical analyses; A.A., R.G., and F.D. worked on the original draft of the manuscript; all authors contributed to the review and editing of the publication.

## Declaration of interests

The authors have no competing interests to declare.

## Notes

### Competing Interest Statement

The authors have declared no competing interest.

### Summary of Updates

The revised manuscript contains clarifications on some of the protocol steps, and lists more items in the key resource table that were missing in the previous version.

